# Circadian clock control of MRTF-SRF pathway suppresses beige adipocyte thermogenic recruitment

**DOI:** 10.1101/2022.04.06.487359

**Authors:** Xuekai Xiong, Weini Li, Ruya Liu, Pradip Saha, Vijay Yechoor, Ke Ma

## Abstract

The morphological transformation of adipogenic progenitors into mature adipocytes requires dissolution of cytoskeleton driven by progressive loss of MRTF-SRF activity. Circadian clock confers temporal control in adipogenic differentiation, and MRTF-SRF inhibits beige adipocyte development. Here we identify that key components of the actin cytoskeleton-MRTF/SRF signaling cascade are direct transcriptional targets of circadian clock, and a clock-MRTF/SRF regulatory axis suppresses beige adipocyte thermogenic recruitment. Genetic loss- or gain-of-functions of the core clock regulator, Brain and Muscle Arnt-like 1 (Bmal1), altered actin cytoskeleton organization, cell shape and SRF activity. Genes involved in actin dynamic-MRTF-SRF pathway display diurnal expression profiles in beige adipose tissue, with identification of Bmal1 direct transcriptional regulation. Using beige adipocyte-selective genetic models together with pharmacological approaches, we demonstrate that *Bmal1* inhibits beige adipogenesis and thermogenic capacity *via* the MRTF-SRF pathway. Selective ablation of *Bmal1* induced beigeing, whereas its targeted overexpression attenuated the beige thermogenic program resulting in obesity. Collectively, our findings define a novel circadian clock control in actin cytoskeleton-MRTF-SRF signaling that suppresses beige adipogenesis to maintain metabolic homeostasis.

## Introduction

The development of mature adipocytes from adipogenic progenitors is characterized by drastic morphological transformation into a lipid-laden spherical shape from fibroblast-like morphology (Kanzaki & Pessin, 2001; Rodriguez Fernandez & Ben-Ze’ev, 1989; Spiegelman & Farmer, 1982). This process requires extensive intracellular actin cytoskeleton remodeling with dissolution of filamentous actin (F-actin) stress fibers, and is known to precede lipid accumulation (Spiegelman & Farmer, 1982). Serum response factor (SRF) and its co-activator, Myocardin-related Transcription Factor-A/B (MRTF-A/B, MKL1/2), mediate the transcriptional response to actin dynamic that alters the ratio between monomeric globular actin (G-actin) and polymerized F-actin (Miralles *et al*, 2003; Olson & Nordheim, 2010; Posern & Treisman, 2006). Besides its established role in driving distinct muscle lineage development (Li *et al*, 2005; Long *et al*, 2007; Wang *et al*, 2002; Wang *et al*, 2004), progressive loss of MRTF/SRF activity is required for cytoskeleton reorganization that enables adipogenic induction during adipocyte formation (Liu *et al*, 2020; Spiegelman & Farmer, 1982). Recent studies indicate that MRTF/SRF-related signaling cascade suppresses mesenchymal precursor commitment and terminal differentiation into beige adipocytes that possess cold-inducible thermogenic capacity, with loss of MRTF-A leading to browning of beige fat depot and improved energy balance (Liu *et al*., 2020; McDonald *et al*, 2015; Rosenwald *et al*, 2017).

Functionally distinct adipose depots, including the classic visceral white adipose tissue and thermogenic brown and beige fat, plays specific roles in energy homeostasis (Cohen & Kajimura, 2021; Sidossis & Kajimura, 2015; Wu *et al*, 2012). While white adipocytes in visceral depots specialize in lipid storage, beige adipocytes residing in subcutaneous fat possess inducible thermogenic activities for energy dissipation. This thermogenic property involves uncoupled mitochondrial respiration mediated by Uncoupling Protein 1 (UCP1) in response to cold exposure or β-adrenergic stimulation. Exploring regulatory pathways to expand or activate the energy-dissipating capacity of thermogenic fat depots may maintain energy balance and protect against obesity and associated metabolic consequences (Boss & Farmer, 2012; Stanford *et al*, 2013; Stanford *et al*, 2015). Functionally divergent adipose depots are derived from distinct mesenchymal precursor lineages with specific molecular pathways governing tissue growth, expansion and functional activation (Cohen & Kajimura, 2021; Sanchez-Gurmaches *et al*, 2016), rendering targeting their metabolic functions *in vivo* challenging (Jeffery *et al*, 2014; Krueger *et al*, 2014; Sanchez-Gurmaches *et al*, 2015). The recent discovery of the inhibitory effect of MRTF/SRF on beige adipogenesis suggests that this regulatory cascade could be targeted to augment beige fat capacity.

The circadian clock is a hierarchical system consisting of the central clock located in the suprachiasmatic nuclei and cell-autonomous peripheral clocks in nearly all tissue and cells (Finger *et al*, 2020; Takahashi, 2017). The central clock receives daily light signal transmitted from retina, which entrains the clock circuits in peripheral tissues under normal physiological conditions. A transcriptional negative feed-back loop, coupled with translational and post-translational regulations, underlies the ∼24 hour rhythmic oscillations in the central and peripheral clock circuits (Takahashi, 2017). Clock transcription activators, Brain and Muscle Arnt-like 1 (Bmal1) and CLOCK (Circadian Locomotor Output Cycles Kaput), form a heterodimer that activates the transcription of their repressors, the Period and Cryptochrome genes. Subsequent Per/Cry-mediated repression of Bmal1/CLOCK activity constitutes the negative feed-back regulation that generates the core clock oscillation. Additional Rev-erbα and ROR-mediated control of *Bmal1* rhythmic expression re-enforces the core clock mechanism. Tissue-intrinsic clock circuits confer temporal control to metabolic pathways to orchestrate metabolic homeostasis (Panda *et al*, 2002), with their disruption leading to the development of insulin resistance and obesity (Scheer *et al*, 2009; Turek *et al*, 2005; Xiong *et al*, 2021). Cell-autonomous peripheral clocks are present in adipose depots (Otway *et al*, 2009; Wu *et al*, 2007; Zvonic *et al*, 2006), and various circadian clock components are known to modulate adipogenesis (Aggarwal *et al*, 2017; Fontaine *et al*, 2003; Grimaldi *et al*, 2010; Guo *et al*, 2012; Kawai *et al*, 2010; Zhu *et al*, 2018). The essential core clock transcription activator Bmal1 inhibits differentiation of white and brown adipocytes (Guo *et al*., 2012; Nam *et al*, 2015b), while its repressor protein Rev-erbα (Nr1d1) promotes brown adipogenesis (Nam *et al*, 2015a).

In response to cell-surface biochemical and extracellular cues, an intricate intracellular signaling cascade stimulates polymerization of monomeric globular actin (G-actin) to form filamentous actin (F-actin) that lowers effective actin monomer concentration (Miralles *et al*., 2003; Olson & Nordheim, 2010). Subsequent release of MRTF-A/B from sequestration by G-actin leads to its nuclear translocation to activate SRF-mediated transcription that drives fundamental cellular behaviors involving cytoskeleton, including adhesion, migration and differentiation in developmental processes. Intriguingly, cyclic circulating cues from serum stimulate actin remodeling in liver and the oscillation of G-actin to F-actin ratio entrains clock, thus connecting actin dynamics with clock modulation (Esnault *et al*, 2014; Gerber *et al*, 2013). Whether the circadian clock exert temporal control on intracellular actin cytoskeleton remodeling is yet to be addressed, and current understanding of regulatory mechanisms modulating MRTF/SRF activity besides actin dynamics is limited. In the current study, we identify that the actin cytoskeleton-MRTF/SRF signaling cascade is subjected to transcriptional control by the circadian clock, and this regulation underlies cell-autonomous clock modulation of beige fat thermogenic capacity.

## Results

### Bmal1 modulates actin cytoskeleton organization and exerts transcriptional control of actin-MRFT/SRF signaling pathway

C3H10T1/2 (10T1/2) are mesenchymal stem cells capable of beige adipocyte lineage commitment (Tseng *et al*, 2008). In these mesenchymal beige precursors with loss- and gain-of-function of *Bmal1* (Nam *et al*., 2015b), we observed striking changes in cell shape with altered actin cytoskeleton organization (Fig. 1A). Inhibition of *Bmal1* by stable shRNA silencing (*shBM*) reduced cell size and the abundance of polymerized F-actin. In contrast, *Bmal1* overexpression (*BMO/E*) augmented actin stress fibers with enlarged size. Global transcriptomics profiling by microarray in *shBM* and scramble control (*shSC)* 10T1/2 cells revealed, in addition to known clock-regulated metabolic and cancer-associated processes, enrichment of differentially-regulated pathways related to cytoskeleton modulation, including focal adhesion, adherent junctions, actin cytoskeleton and cytokine-receptor interaction (Fig. 1B & Fig. S1), RT-qPCR analysis validated that various components involved in cytoskeleton-MRTF/SRF signaling cascade were down-regulated with silencing of *Bmal1. Srf* transcript was reduced to ∼50% of *shSC* (Fig. 1C). Expression of known MRTF-SRF target genes, *vinculin* (*Vcl), connective tissue growth factor* (*Ctgf)* and *Four and half LIM domain protein 1 and 2* (*Fhl1 & Fhl2)*, were nearly abolished, suggesting attenuated SRF activity. SRF protein was reduced in cells with *Bmal1* silencing in the presence or absence of serum stimulation (Fig. 1D), whereas ectopic expression of *Bmal1* increased SRF protein level (Fig. 1E). Analysis of SRF activity by an SRF response element (SRE)-driven luciferase reporter (SRE-Luc) revealed *Bmal1* modulation of SRF-mediated transcription, with *Bmal1* knockdown significantly attenuating (Fig. 1F) and its overexpression augmenting luciferase activity (Fig. 1G). MRTF/SRF activity was progressively suppressed during adipogenic differentiation (Liu *et al*., 2020). In *Bmal1*-deficient 10T1/2 cells at day 9 of beige differentiation, actin filament was nearly completely lost with attenuated vinculin staining, whereas F-actin and vinculin-stained focal adhesion were largely maintained *Bmal1*-overexpressing cells (Fig. 1H). Prominent loss of F-actin organization and focal adhesion contact by vinculin staining was observed throughout the beige differentiation time course in *shBM* as compared to *shSC* controls (Suppl. Fig. S2A & S2B).

**Figure 1.**
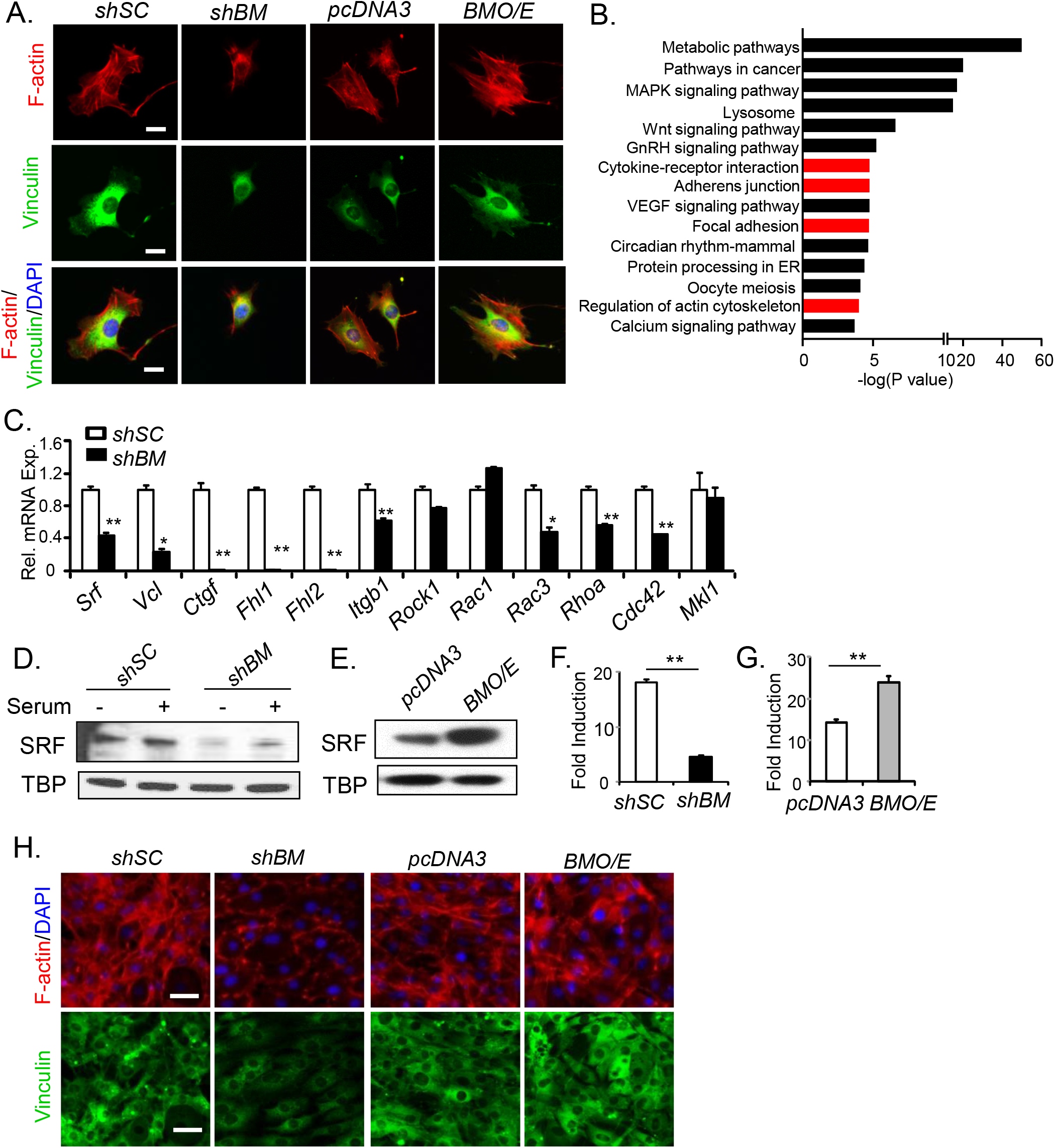
Bmal1 regulation of the actin cytoskeleton and MRTF/SRF signaling pathway. (A). Representative confocal images of F-actin staining by phalloidin (red) and vinculin (green) immunostaining in C3H10T1/2 cells containing stable expression of shRNA of scrambled control (*shSC*) and Bmal1 (*shBM*), or *pcDNA3* and Bmal1 cDNA (*BMO/E)*. Scale bar: 25 µm.(B) KEGG analysis of enriched pathways of differentially expressed genes in *shBM* vs. *shSC* 10T1/2 cells (n=3). (C) RT-qPCR analysis of cytoskeleton and SRF-related genes in *shSC* or *shBM* 10T1/2 cells (n=4). (D, E) Immunoblot analysis of SRF with or without serum stimulation in shSC and *shBM* (D), or *pcDNA3* and *BMO/E* cells (E). (F & G) SRE-Luc luciferase reporter assay of SRF activity in response to serum in *shSC* and *shBM* (F). or *pcDNA3* and *BMO/E* cells (G). Values were expressed as fold induction of serum stimulation over basal control without serum. **: P≤0.01 vs. respective controls (n=4). (H). Representative images of F-actin and vinculin staining in *shSC* and *shBM*, or *pcDNA* and *BMO/E* cells at day 9 of 10T1/2 beige adipogenic differentiation. Scale bar: 100 µm.

Based on these findings, we tested whether Bmal1 exert direct transcriptional control of genes within the actin cytoskeleton-MRTF/SRF signaling cascade. Screening for putative Bmal1 binding sites, the E- or E’-box elements (Chatterjee *et al*, 2013; Rey *et al*, 2011), in gene cis-regulatory regions using TRANSFAC, identified several candidate targets within this pathway. ChIP-qPCR analysis revealed enriched Bmal1 chromatin occupancies of proximal promoters of genes involved in cytoskeleton-SRF regulation, including *Srf, Vcl, Ctgf, Fhl1* and *Fhl2* (Fig. 2A). Comparing to the E-box site on a known Bmal1 target, *Rev-erbα* promoter, these enrichments were less robust (Fig. 2A). In *Bmal1*-deficient 10T1/2 cells, Bmal1 occupancy of these chromatin regions were largely abolished to a similar degree as IgG control. To determine whether Bmal1 transcriptional regulation confers circadian oscillation of the identified targets involved in cytoskeleton dynamics, we induced synchronization through serum shock and analyzed gene expression profiles across two circadian cycles (Chatterjee *et al*., 2013). As expected, serum shock induced *Bmal1* rhythm that was significantly dampened in *shBM* cells (Fig. 2B). Similar serum shock-induced oscillatory profiles, together with attenuation by *Bmal1* inhibition were observed in *Srf* and MRTF-SRF target genes, *Vcl* and *Fhl*. In inguinal subcutaneous white adipose tissue (iWAT), a representative beige adipose depot, both SRF and MRTF-A protein displayed daily rhythmicity with peak levels detected at early morning time point that were in phase with Bmal1 expression (Fig. 2C & 2D).

**Figure 2.**
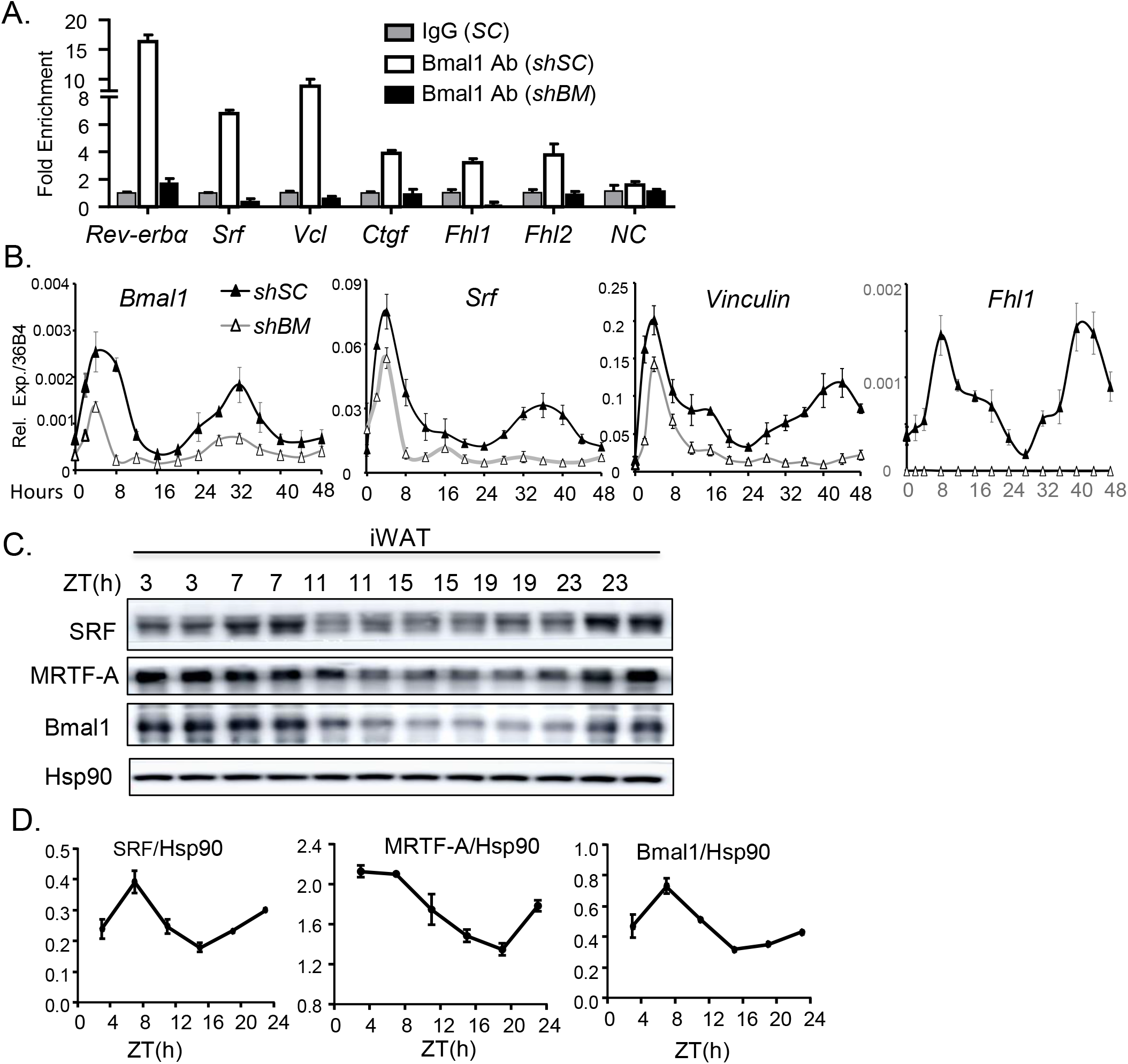
Bmal1 exerts direct transcriptional control of the cytoskeleton-MRTF/SRF pathway. (A) ChIP-qPCR analysis of Bmal1 chromatin occupancy on regulatory regions of MRTF/SRF pathway genes in *shSC* and *shBM* C3H10T1/2 cells. Values were calculated as % of input and normalized to IgG controls (n=4). NC: negative control of *Rev-erba* promoter upstream primers. (B) RT-qPCR analysis of circadian expression induced by serum shock every four hours for 48 hours in *shSC* and *shBM* 10T1/2 cells (n=3). (C, D) Immunoblot analysis (C) with quantitation (D) of circadian expression of SRF and MRTF-A protein in iWAT of 10-12 week-old female mice for 24 hours. ZT, Zeitgieber time. Pooled samples of n=4-5/time point.

### Selective ablation of Bmal1 in beige adipose tissue promotes browning and improves metabolic homeostasis

Prx1-Cre selectively targets subcutaneous beige adipose tissue progenitors and mature adipocytes without visceral white adipose or interscapular brown adipose tissue involvement (Jeffery *et al*., 2014; Krueger *et al*., 2014; Sanchez-Gurmaches *et al*., 2015). To determine the role of the clock-actin-MRTF/SRF regulation in beige adipocyte development *in vivo*, we used Prx1-Cre transgenic mice, *via* a cross with floxed *Bmal1* allele (BM^fl/fl^), to achieve *Bmal1* ablation in beige fat. In Prx1-Cre^+^/BM^fl/fl^ (*bBMKO*) mice as compared to Prx1-Cre^-^/BM^fl/fl^ (*bBMCtr*) littermate controls, absence of Bmal1 protein was confined to beige iWAT depot but not in gonadal/visceral white adipose tissue (gWAT), BAT or liver (Fig. 3A). Loss of *Bmal1* transcript was corroborated by RT-qPCR analysis in iWAT, with attenuated expression of Bmal1 target gene, *Rev-erbα* (Fig. 3B). Notably, compared to *bBMCtr*, 6-week-old *bBMKO* mice iWAT displayed significant beigeing with mitochondrial-rich cytoplasmic staining and multi-locular lipid droplets (Fig. 3C, upper panel). Ucp-1 protein level in *Bmal1-*deficeint iWAT was elevated as shown by immunostaining (Fig. 3C, lower panel). Consistent with the beigeing phenotype, RNA-seq profiling revealed up-regulation of mitochondria-related processes in *bBMKO* iWAT (Fig. 3D). In addition, KEGG analysis identified fatty acid and lipid metabolism were among the top enriched pathways (Fig. 3E), ad these processes overlapped with the metabolic signature induced by cold acclimation (Fig. 3F), suggesting a metabolic phenotype resembling cold-induced thermogenic induction in the *Bmal1-*deficient beige fat. Indeed, thermogenic genes were markedly induced in *bBMKO* iWAT, with ∼4-6 fold up-regulation of *Ucp-1, Cebpb* and *Pparγ* (Fig. 3G). UCP-1 protein was elevated in *Bmal1*-deficient iWAT under ambient temperature, and further induced upon cold stimulation (Fig. 3H). PGC-1α was moderately increased in *bBMKO*, while C/EBPα and PPARγ protein levels trended toward lower as compared to controls.

**Figure 3.**
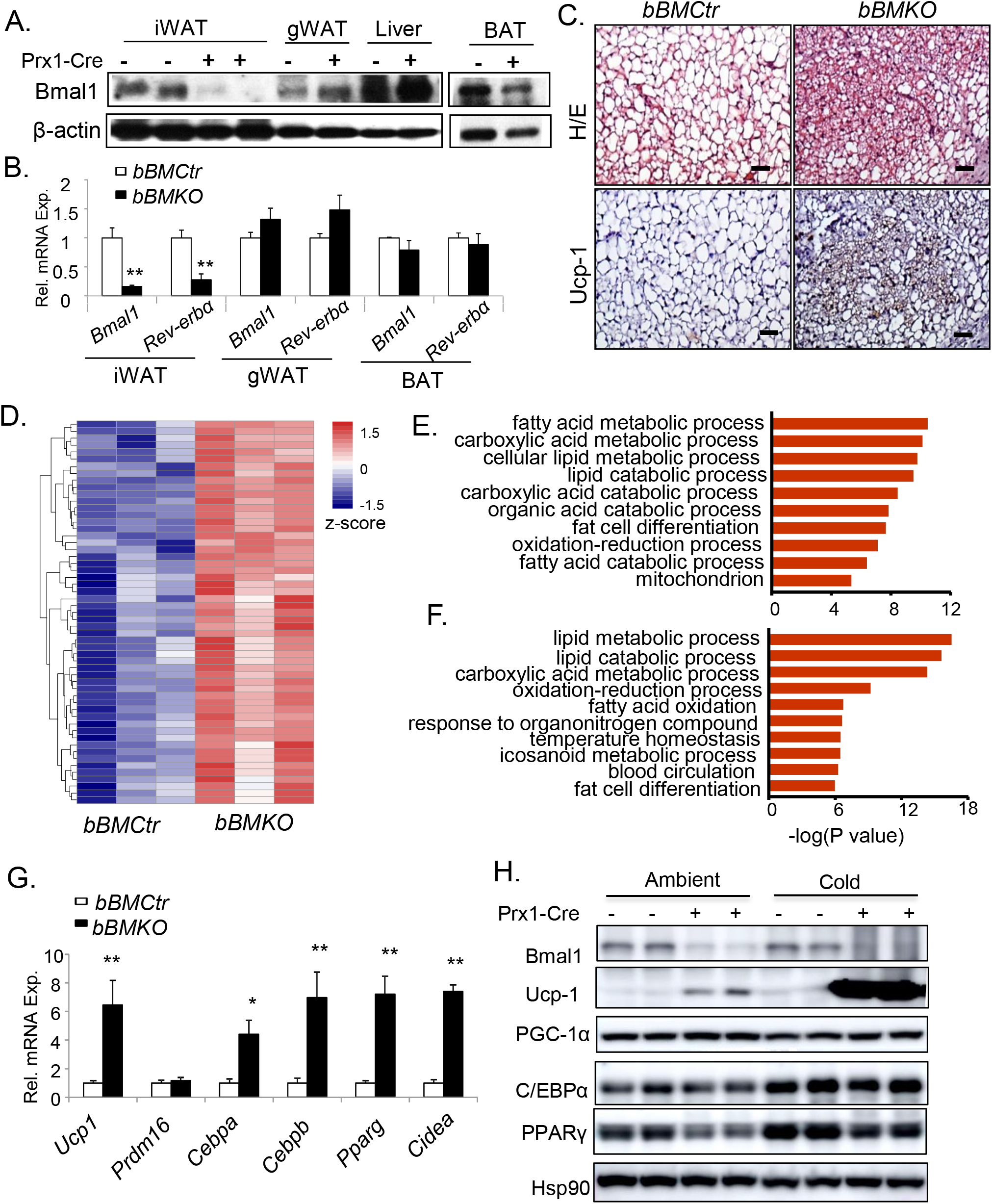
Selective ablation of *Bmal1* induces browning of beige fat. (A, B) Immunoblot analysis of Bmal1 protein (A), and mRNA level (B) in fat depots and the liver of *bBMCtr* and *bBMKO* male mice. iWAT: inguinal subcutaneous white adipose tissue; gWAT: gonadal visceral white adipose tissue; BAT: brown adipose tissue. (C) Representative images of H&E histology (upper panels) with UCP-1 immunostaining (lower panels) of iWAT in 6-week-old *bBMCtr* and *bBMKO* male mice. (D) Heatmap representation of up-regulated genes in mitochondria-related processes in iWAT from male *bBMKO* mice as compared to *bBMCtr* by RNA-seq analysis. (E & F) KEGG analysis of up-regulated pathways in b*BMKO* iWAT as compared to *bBMCtr* mice in ambient condition (E), and cold acclimation-induced pathways as compared to ambient condition in *bBMCtr* mice (F). N=3/group. (G) RT-qPCR analysis of thermogenic and adipogenic genes in *bBMKO* and *bBMCtr* mice under ambient condition (n=5/group). (H) Immunoblot analysis of thermogenic gene expression in iWAT of female *bBMCtr and bBMKO* mice under ambient or cold acclimation (n=pooled sample of 3-4 mice/lane).

Given the metabolic benefit of inducible beige fat thermogenic activity, we examined whether *Bmal1* ablation*-*induced beigeing of subcutaneous fat depot is sufficient to promote energy balance. Loss of *Bmal1* in beige fat did not alter body weight or total fat mass in 8-week-old mice, as assessed by NMR (Fig. S3A& S3B). By 6 months of age, however, *bBMKO* mice showed significant reduction of body weight with ∼40% lower fat mass, as compared to littermate controls (Fig. 4A). Both iWAT and gWAT weight were decreased, but not BAT (Fig. 4B). *Bmal1*-deficient iWAT maintained enhanced beige characteristic at this age as compared to age-matched *bBMCtr* mice, and adipocytes were smaller in iWAT and gWAT (Fig. 4C). The age-dependent reduction of adiposity with beige-selective *Bmal1* ablation suggests potential cumulative effects of beigeing on energy balance. *bBMKO* mice at ambient temperature displayed a tendency toward higher oxygen consumption, which became significantly elevated than controls following cold acclimation at 12°C (Fig. 4D & 4E). Marked induction of lipid catabolic process in *Bmal1*-deficient iWAT at ambient temperature with stimulation by cold from RNA-seq analysis further corroborated these findings (Fig. 4F).

**Figure 4.**
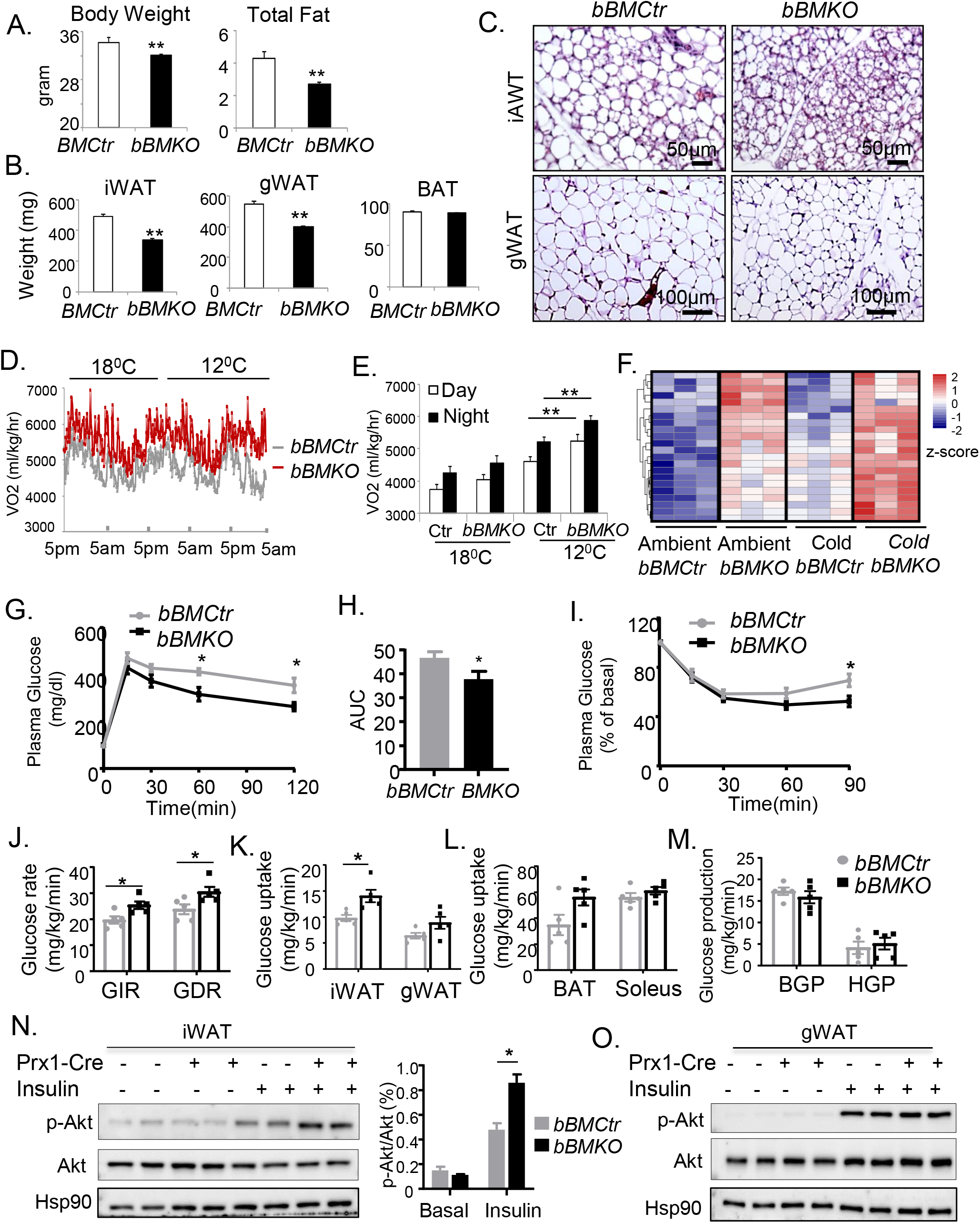
*Bmal1* deficiency in beige fat augments energy expenditure with improved glucose metabolism. (A-C) Analysis of total body weight and fat mass by NMR (A), fat tissue weight (B), and representative H&E histology of iWAT and gWAT (C) in 6-month-old male *bBMKO* and *bBMCtr* mice. N=6-8/group. (D, E) Metabolic cage analysis of oxygen consumption rate (VO2) in male *bBMKO* and *bBMCtr* mice as shown by average tracing (D), and quantification (E) under ambient temperature and cold acclimation at 18°C or 12°C. (n=6/group). (F) Heatmap representation of up-regulation of fatty acid metabolic pathway by RNA-seq analysis in female *bBMKO* and *bBMCtr* mice under ambient or cold conditions (n=3/group). (G-I) Glucose tolerance test (GTT, G) with area under the curve (AUC) quantification (H), and (I) insulin tolerance test of 10-12 week-old male *bBMCtr* and *bBMKO* mice at ambient temperature (n=6-7/group). (J-M) Hyperinsulinemic-euglycemic clamp analysis of steady-state glucose infusion rate (GIR) and glucose disposal rate (GDR) (J), quantification of glucose uptake in iWAT, gWAT (K), BAT and Soleus muscle (L), and hepatic glucose production rate (HGP, M) from male *bBMKO* and *bBMCtr* mice (n=5/group). BGP: basal glucose production. (N & O) Akt phosphorylation in response to intraperitoneal 0.5U/Kg insulin stimulation in iWAT (N), and gWAT (O) in female *bBMKO* and *bBMCt*r mice (n=3 pooled mice/lane). *, **: P≤0.05 or 0.01 by Student’s t test.

To determine whether beigeing in *bBMKO* mice improves glucose homeostasis, we performed glucose tolerance test and found lower blood glucose levels at 60 and 120 minutes following intraperitoneal glucose challenge in *bBMKO* mice (Fig. 4G), with significant reduction of area under the curve (Fig. 4H). *bBMKO* mice also displayed augmented insulin sensitivity as shown by insulin-induced glucose lowering effect (Fig. 4I). We further examined glucose disposal in these mice under a steady-state condition during hyperinsulinemic-euglycemic insulin clamp. Both glucose infusion and disposal rates were significantly elevated in *bBMKO* mice as compared to *bBMCtr* (Fig. 4J). Enhanced glucose disposal was likely attributed to glucose uptake in *Bmal1*-deficient iWAT (Fig. 4K), while its entry into gWAT, BAT or soleus muscle were comparable to that of the controls (Fig. 4K & 4L). Hepatic glucose production at basal or clamped state were not significantly altered (Fig. 4M). Direct examination of insulin signaling revealed enhanced Akt phosphorylation in *Bmal1*-deficient iWAT upon stimulation, indicating improved beige fat insulin sensitivity (Fig. 4N). In contrast, Akt activation in gWAT was similar between *bBMKO* and *bBMCtr* mice (Fig. 4O). Given the improved energy balance with Bmal1 ablation in beige fat, we determined *bBMKO* mice response to high-fat diet (HFD) challenge. When subjected to HFD for 14 weeks, these mice maintained lower body weight than littermate controls (Fig. 5A). iWAT and gWAT from HFD-fed *bBMKO* mice displayed reduced adipocyte hypertrophy as compared to *bBMCtr* (Fig. 5B), with a marked shift toward smaller adipocytes as revealed by size distribution (Fig. 5C & 5D). iWAT and gWAT fat pad weight and combined mass were significantly lower in *bBMKO* mice (Fig. 5E). Moreover, *bBMKO* mice displayed augmented insulin sensitivity on HFD, as indicated by stronger glucose-lowering effect stimulated by insulin as compared to controls (Fig. 5F & 5G).

**Figure 5.**
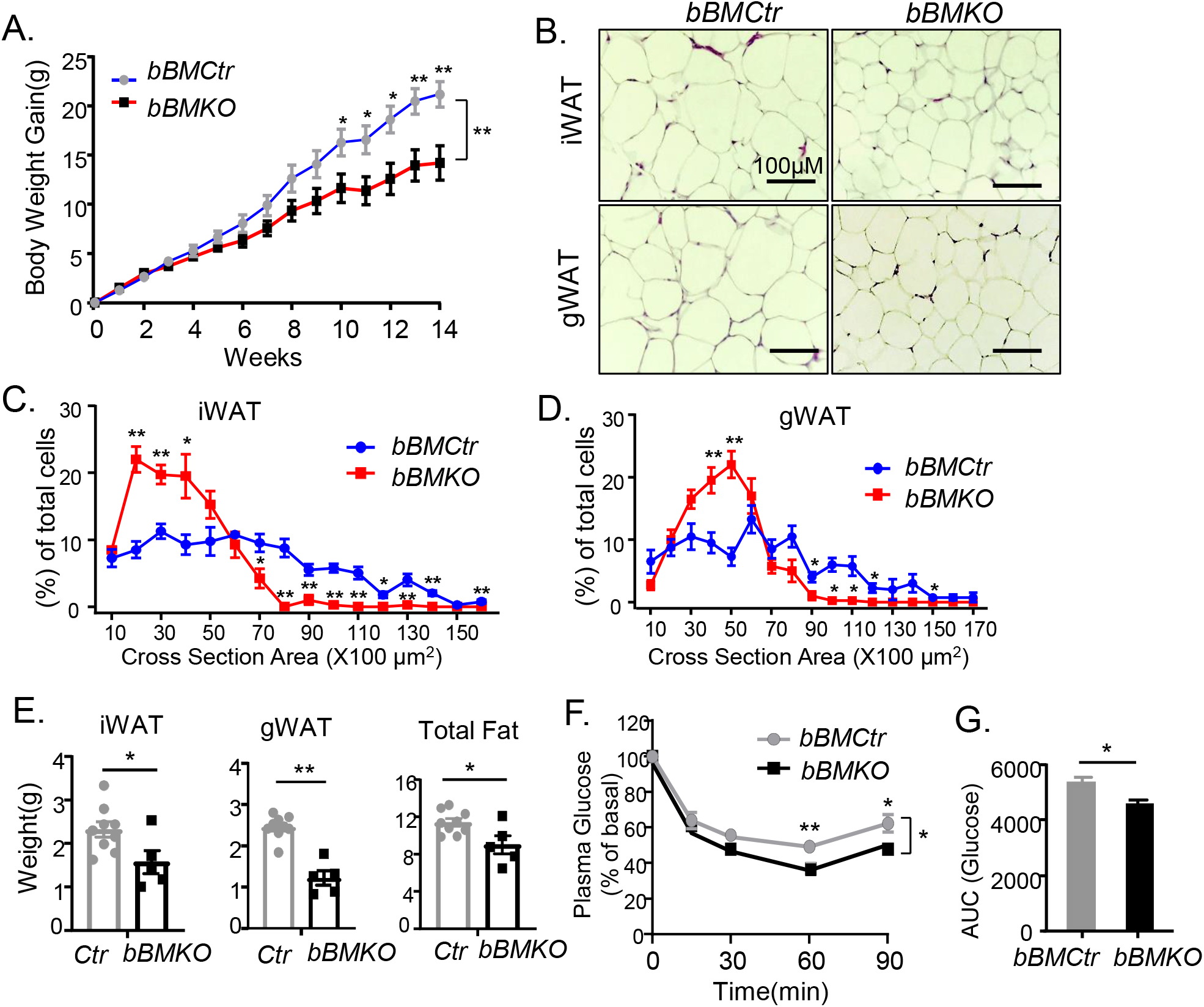
*Bmal1* deficiency in beige fat protects against HFD-induced obesity. (A) Body weight growth curve as shown by weekly weight gain of male *bBMCtr* (n=9) and *bBMKO* (n=5) mice fed 45% high fat diet for 14 weeks. *, **: P≤0.05 or 0.01 one-way ANOVA with post-hoc pairwise t test. (B-D) H&E histology of iWAT and gWAT after high fat diet (B), with quantification of adipocyte size distribution (C & D). (E) Tissue weight analysis of iWAT, gWAT and total fat mass in high fat diet-fed *bBMCtr* (n=9) and *bBMKO* mice (n=5). (F & G) Plasma glucose levels during insulin tolerance test (F), and area under the curve quantification (G) in male *bBMCtr* (n=6) and *bBMKO* mice (n=5) after high fat diet feeding. *, **: P≤0.05 or 0.01 by Student’s *t* test or ANOVA with post-hoc pairwise t test.

### Beige fat Bmal1 deficiency in vivo dampens MRTF-SRF signaling with enhanced beige preadipocyte differentiation

In *Bmal1*-deficient iWAT, KEGG analysis revealed extracellular matrix (ECM) and cell adhesion-related pathways as among the top down-regulated processes (Fig. 6A), and they overlapped with pathways suppressed by cold acclimation (Fig. 6B). Notably, *Bmal1* deficiency and cold-induced suppression of ECM-related processes in beige depot were synergistic (Fig. 6C). Ingenuity Pathway Analysis (IPA) of differentially-expressed genes (DEGs) identified SRF and MRTF-A as up-stream regulators (Fig. S4A), with down regulation of MRTF-SRF positive transcriptional targets (Fig. S4B). *Srf* and *Mkl1* (*Mrtf-a*) expression were moderately reduced in *Bmal1*-dificient iWAT at ambient temperature and both transcripts were markedly suppressed by cold, while cold stimulation alone did not affect *Srf* or *Mkl1* (Fig. 6D). As Prx1-Cre-mediated gene ablation targets both beige progenitors and mature adipocytes, we determined how Bmal1 modulates cytoskeleton-MRTF/SRF cascade in the stomal vascular fraction (SVF) containing preadipocytes or mature adipocytes isolated from iWAT beige depot. Significant down-regulations of *Srf* and *Vcl* were confined to mature beige adipocytes only in Bmal1-deficient iWAT (Fig. 6E), but not the SVFs. Interestingly, *Srf* was induced in adipocytes as compared to SVF in *bBMCtr*, although *Mkl1* and *Mkl2* were suppressed together with reduced *Ctgf* expression suggesting attenuated MRTF/SRF transcription activity. Corroborating observed beigeing in *bBMKO* iWAT, the thermogenic program in isolated *Bmal1*-deficient mature beige adipocytes, including *Ucp-1, Dio2* and *Pgc-1a*, were induced (Fig. 6F), whereas adipogenic gene expression were comparable to that of controls (Fig. 6G).

**Figure 6.**
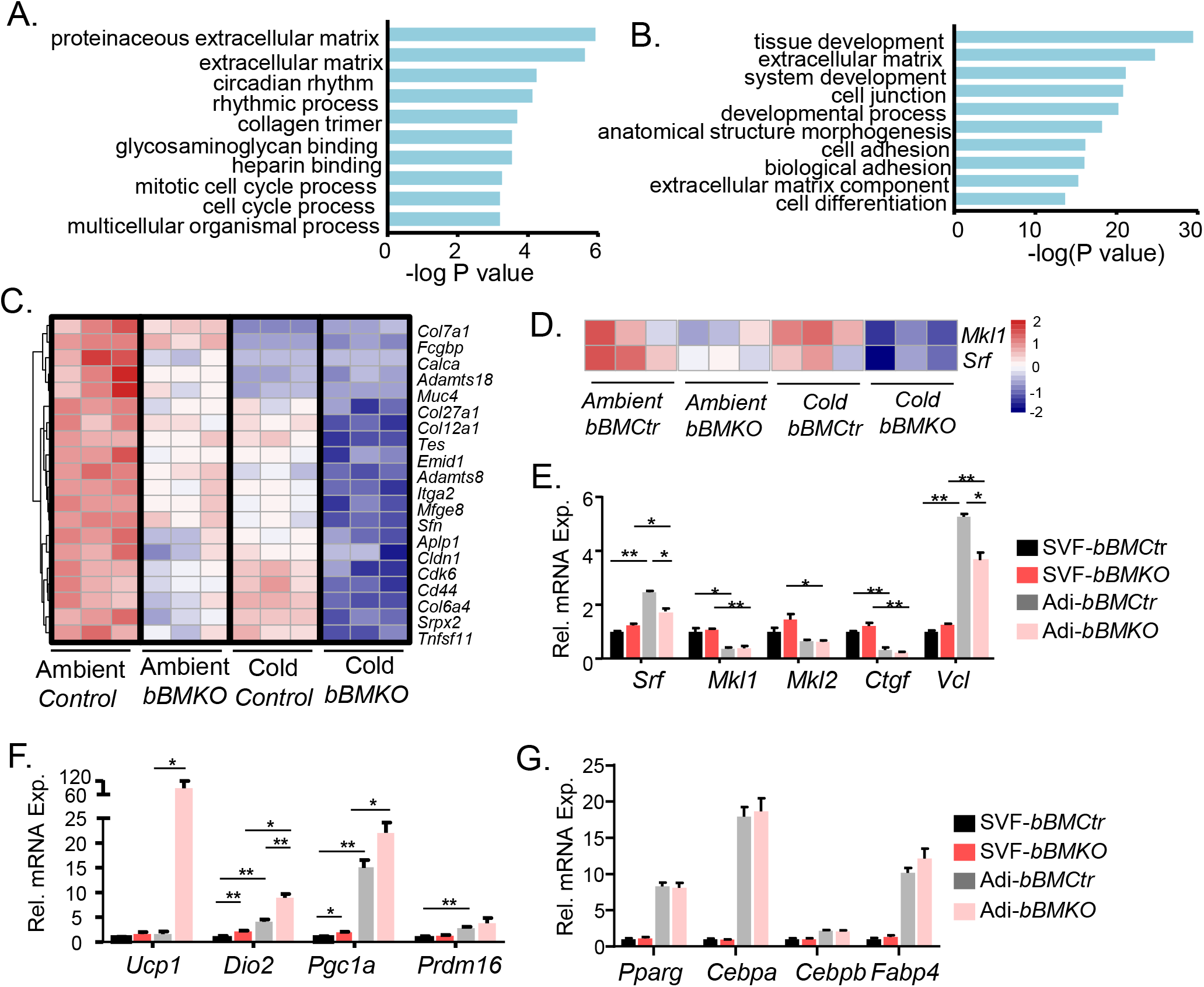
Loss of *Bmal1* in beige fat suppresses MRTF-SRF pathway. (A, B) KEGG analysis of RNA-seq of down-regulated pathways in iWAT from *bBMKO* and *bBMCtr mice* (A), and overlap with cold acclimation-suppressed processes in *bBMCtr mice* (B, n=3/genotype). (C, D) Heatmap representation of differentially expressed genes in extracellular matrix and cell adhesion pathways (C), and expression of *Mkl1* and *Srf* by RNA-seq (D) in iWAT from female *bBMCtr* and *bBMKO* mice under ambient condition or cold acclimation. (E-G) RT-qPCR analysis of genes related to MRTF-SRF pathway (E), thermogenic (F), and adipogenic program (G) in isolated stromal vascular fraction (SVF) or mature adipocytes (Adi) from iWAT of *bBMKO* and *bBMCtr* mice (n=5-6/group). *, **: P≤0.05 or 0.01 by Student’s t test.

Transcriptomic profiling in *bBMKO* iWAT revealed marked up-regulation of fat cell differentiation pathway (Fig. S4C). We thus tested whether impaired MRTF-SRF activity, as a result of loss of *Bmal1* regulation, underlies the beigeing phenotype *in vivo* by promoting beige adipogenic precursor differentiation. Upon beige induction for 4 and 6 days, *Bmal1*-deficient SVF displayed more abundant lipid accumulation with enhanced mitochondrial staining than *bBMCtr* cells (Fig. 7A & 7B). Thermogenic genes, including *Ucp-1, Pgc1α* and *Diodinase 2* (*Dio2*), were induced in *Bmal1*-deficient beige SVFs (Fig. 7C), together with elevated expression of the early brown lineage marker *Prdm16*. Genes involved in mitochondrial fatty acid transport and oxidation were also induced (Fig. S4D). In comparison, adipogenic gene expression were mostly comparable to that of controls (Fig. 7D), similar to the finding from isolated adipocytes. To determine whether MRTF-SRF modulation underlies enhanced *bBMKO* beige progenitor differentiation, we treated these cells with Jasplakinolide, an actin polymerizing agent that induces MRTF co-activation of SRF (Allingham *et al*, 2006). Jasplakinolide strongly inhibited differentiation of normal control cells, as expected with activation of MRTF/SRF. Notably, Jasplakinolide was able to suppress beige differentiation of *Bmal1*-deficient SVF, suggesting that activating MRTF/SRF can rescue the effect loss of *Bmal1* on beige adipogenesis (Fig. 7E & 7F).

**Figure 7.**
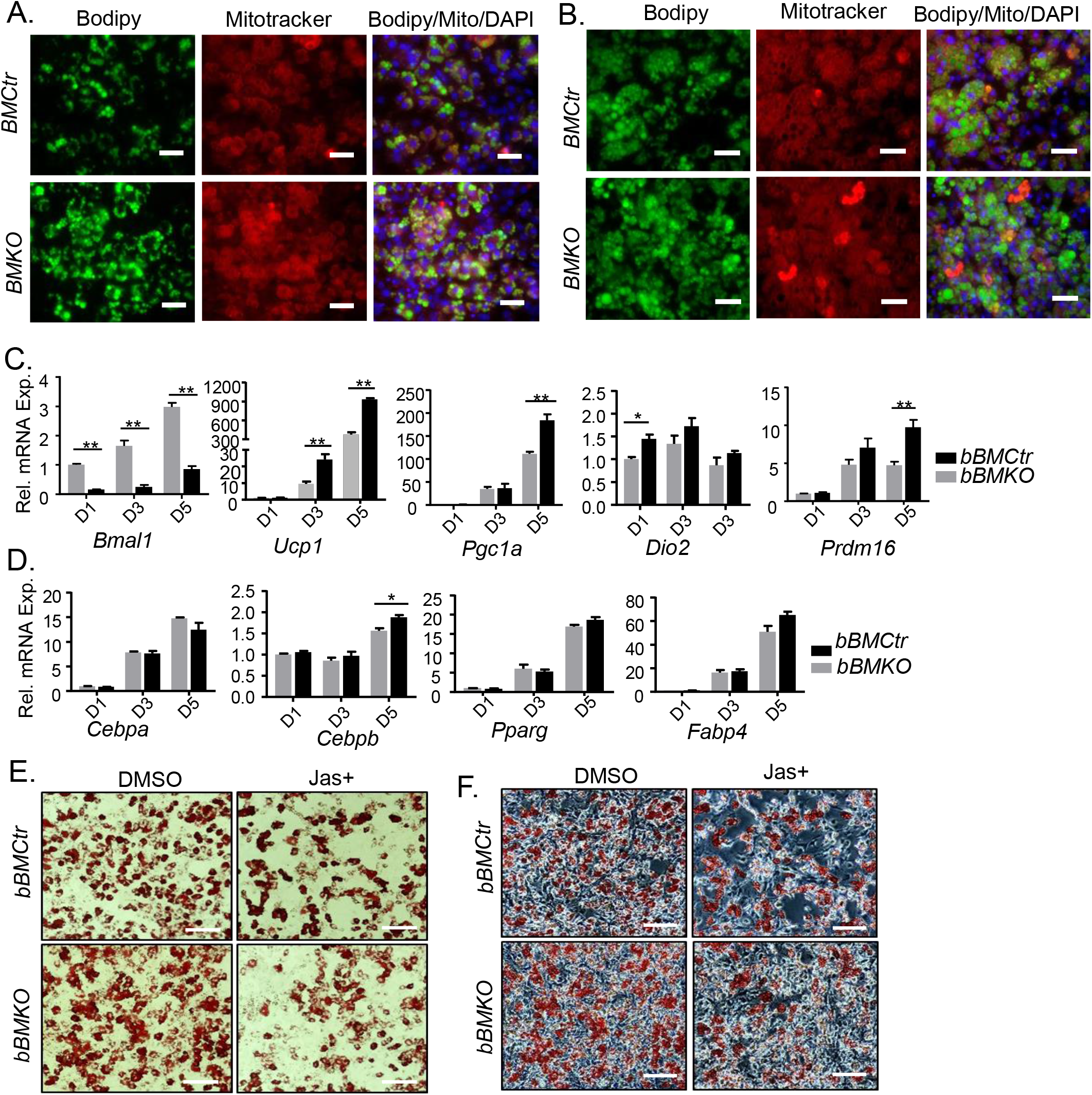
Loss of *Bmal1* promotes beige preadipocytes differentiation. (A, B) Representative images of Bodipy (green) and Mitotracker Red staining at day 4 (A) and day 6 (B) of beige adipogenic differentiation using SVF isolated from iWAT of *bBMCtr* and *bBMKO* mice. (n=5 mice pooled/group). Scale bar: 50 μm. (C, D) RT-qPCR analysis of gene expression during beige adipogenic differentiation time course at day 1, 3 and 5 of adipogenic markers (C) and thermogenic gene program (D). (n=3/group). (E, F) Representative images of oil-Red-O staining (E)with phase-contrast images (F) of day 5-differentiated beige adipocytes from iWAT SVF of *bBMKO* and *bBMCtr* mice with or without Jasplakinolide (Jas+, 0.1 μM) through differentiation. Scale bar: 100 μm.

### Bmal1 overexpression in beige fat enhances MRTF-SRF signaling with impaired beigeing

To further determine whether a Bmal1-cytoskeleton-MRTF/SRF regulatory axis is sufficient to impact beige thermogenic capacity, we generated a conditional *Bmal1* knock-in model *via* CRISPR-mediated gene targeting in the ROSA26 locus. The targeting allele consisted of a CMV promoter-driven Flag-tagged *Bmal1* with bi-cistronic GFP following a Lox-Stop-Lox cassette (Fig. S4A). Floxed knock-in (*bKICtr*) mice were crossed with *Prx1*-Cre to achieve beige fat-selective *Bmal1* overexpression (*bBMKI*), with resulting tagged knock-in Bmal1 protein expression (∼95 kD) detected at ∼5-6 times higher than the endogenous level (∼80 kD) along with GFP (Fig. 8A). Knock-in expression was not present in BAT, liver or skeletal muscle, and weak level was detected in gWAT. Consistent with Bmal1 transcriptional control of SRF, *bBMKI* iWAT displayed elevated SRF protein but not gWAT (Fig. 8B). SRF target genes, *Vcl* and *Fhl1*, were induced in *Bmal1*-overexpressing beige fat, suggesting augmented MRTF/SRF activity, while *Ucp-1* and *Dio2* were suppressed (Fig. 8C). 6-week-old *bBMKI* beige depot lost beige characteristics observed in controls with marked adipocyte hypertrophy, with the histology resembling that of visceral fat (Fig. 8D). Visceral fat in *bBMKI* mice also displayed a tendency toward larger adipocytes. When subjected to cold, *bBMKI* iWAT lacked robust inductions of Ucp-1 and PPARγ found in age-matched *bKICtr* (Fig. 8E). In isolated beige adipocytes from *bBMKI* mice, *Srf* and *Vcl* were significantly up-regulated, while their expression were comparable to controls within the SVF fraction (Fig. 8F). To test whether increased MRTF/SRF activity underlies impaired beigeing in *bBMKI* beige adipocytes, SVF isolated from *bBMKI* were treated with CCG-1423, a specific inhibitor of MRTF/SRF-mediated transcription (Evelyn *et al*, 2010; Evelyn *et al*, 2007). *Bmal1* overexpression suppressed beige adipogenic differentiation of *bBMKI* SVF, as expected with enhanced MRTF/SRF signaling (Fig. 8G). CCG-1423 promoted beige differentiation of *KICtr* cells, as indicated by Bodipy and mitotracker staining. Notably, CCG-1423 induction of beige adipogenesis was abolished in Bmal1-overexpressing SVFs, suggesting that *Bmal1* gain-of-function confers resistance to MRTF/SRF inhibition.

**Figure 8.**
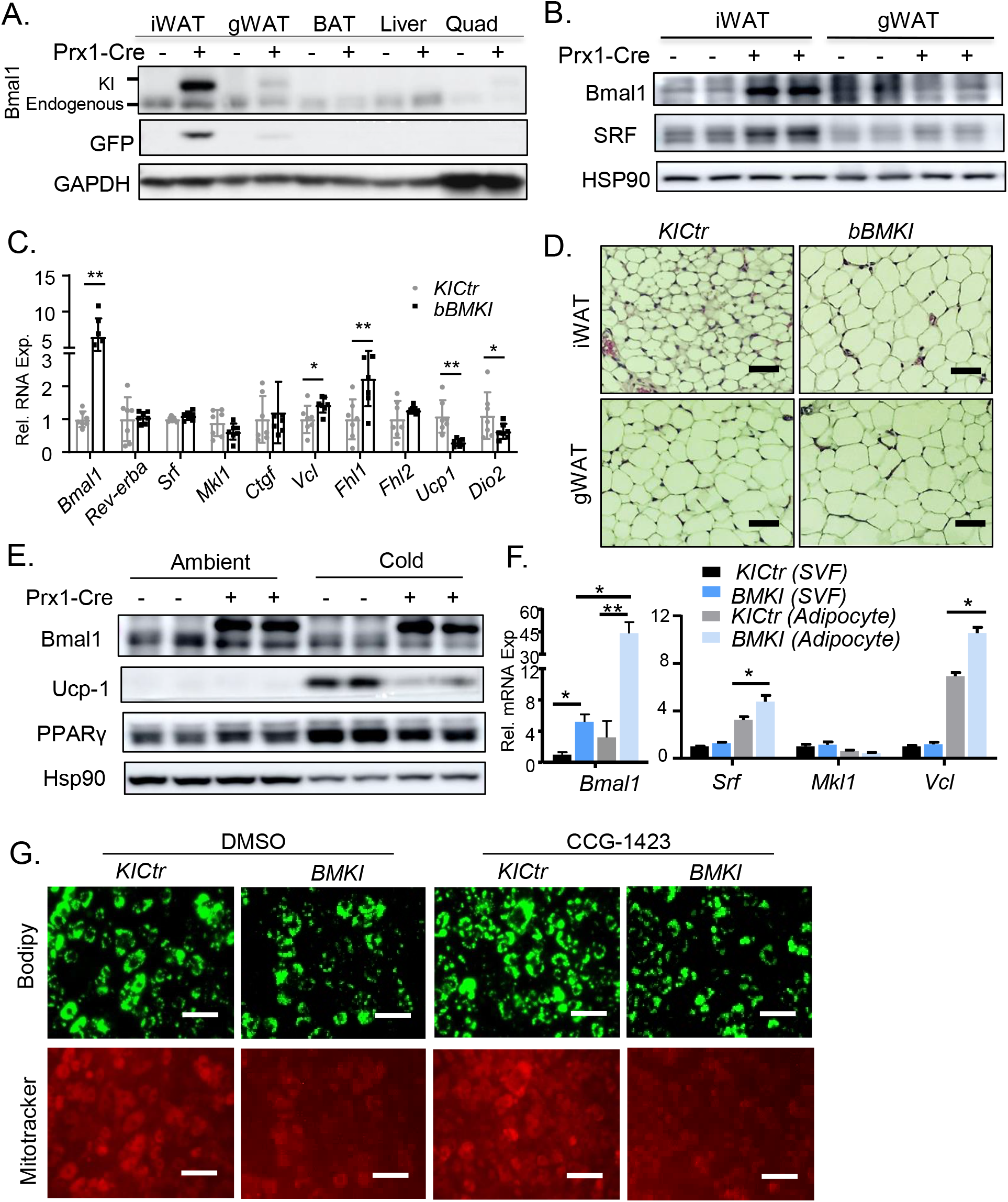
Beige fat-selective *Bmal1* overexpression promotes SRF pathway and suppresses thermogenic gene program. (A) Immunoblot analysis of knock-in (∼95 kD) and endogenous *Bmal1* (∼80 kD) protein in female *bKICtr* and *bBMKI* mice. (Pooled samples of n=4-5 mice/lane). (B) Immunoblot analysis of Bmal1 and SRF in iWAT and gWAT depots in female *bKICtr* and *bBMKI* mice (pooled sample of n=3 mice/lane). (C) RT-qPCR analysis of MRTF/SRF pathway-related and thermogenic gene expression in iWAT of male *bKICtr* (n=6) and *bBMKI* mice (n=7). (D) Representative H&E histology of iWAT and gWAT from 6-week-old male *KICtr* and *bBMKI* mice. Scale bar: 50 μm. (E) Immunoblot analysis of thermogenic and adipogenic factors in *bKICtr* and *bBMKI* mice at ambient temperature or after cold-acclimation. Pooled sample of n=4-5 mice/group. (F) RT-qPCR analysis of MRTF/SRF pathway genes in isolated stromal vascular fraction (SVF) or mature adipocytes from iWAT of *bKICtr* and *bBMKI* mice (n=5-6/group). (G) Representative images of Bodipy (green) and Mitotracker staining (red) at day 5 of beige differentiation of *bKICtr* and *bBMKI* SVF treated with CCG-1423 (5 μM) or DMSO. Scale bar: 70 μm.

Mice overexpressing *Bmal1* in beige fat demonstrated an age-dependent obese phenotype. iWAT and gWAT in three-month-old *bBMKI* displayed striking adipocyte hypertrophy (Fig. 9A), with marked shift of size distribution toward larger adipocytes as compared to age-matched controls. Adipocyte hypertrophy in these depots were reflected by ∼30% increase in respective iWAT and gWAT weight (Fig. 9B & 9C), resulting in increased total fat mass (Fig. 9D) and body weight (Fig. 9E). *bBMKI* BAT displayed enhanced lipid accumulation (Fig. S5B) with increased tissue weight (Fig. S5C). Analysis of energy balance revealed elevated oxygen consumption in *bBMKI* mice confined to the dark active phase (Fig. S5D), together with elevated activity level (Fig. S5E) without altering food intake ((Fig. S5F). The loss of beigeing with resultant obesity led to impaired insulin sensitivity in *bBMKI* mice, as shown by attenuated glucose-lowering response to insulin (Fig. 9F), and the impairment of insulin response was more pronounced following cold acclimation (Fig. 9G). Furthermore, in both subcutaneous beige and visceral fat depots, insulin stimulation of Akt phosphorylation was reduced in *bBMKI* as compared to KI controls (Fig. 9H & 9I). Taken together, *Bmal1* gain-of-function in beige fat depot is sufficient to suppress beigeing that negatively impacts whole-body metabolic regulation.

**Figure 9.**
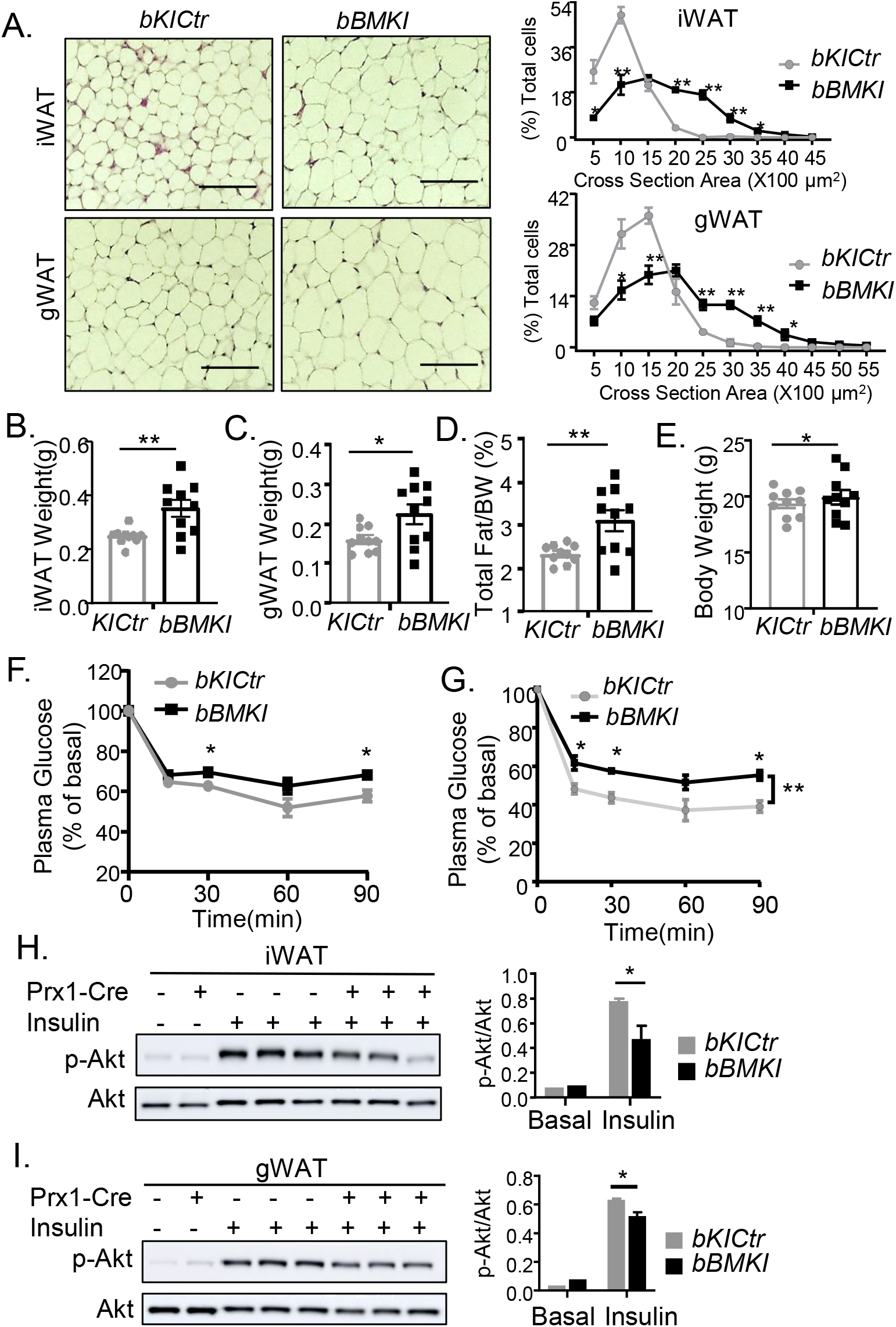
*Bmal1* overexpression in beige fat induces obesity and insulin resistance. (A) H&E histology of iWAT and gWAT from 3-month-old male *KICtr* and *bBMKI* mice with corresponding quantification of adipocyte size distribution. Scale bar: 100 μm. (B-E) Tissue weight analysis of iWAT (B), gWAT (C), total fat mass (D) and body weight (E) in 3-month-old *KICtr* and *bBMK*I mice (n=10/group). (F & G) Plasma glucose levels during insulin tolerance test (0.75U/Kg) in *KICtr* and *bBMKI* mice under ambient temperature (F), or after cold acclimation at 100C for 2 weeks (G). N=6-7 mice/group. *, **: P≤0.05 or 0.01 by one-way ANOVA with post-hoc pairwise t test. (H & I) Immunoblot analysis of Akt phosphorylation in response to 0.5U/Kg insulin stimulation in iWAT (H), or gWAT (I) from female *bKICtr* and *bBMKI* mice with quantification. Pooled samples of n=4-5 for basal and n=3 for insulin-stimulated groups. *: P≤0.05 by student’s t test.

## Discussion

Actin cytoskeleton and modulation of MRTF/SRF activity are fundamental mechanisms underlying stem cell lineage commitment and developmental processes (Connelly *et al*, 2010; McBeath *et al*, 2004; Olson & Nordheim, 2010; Pipes *et al*, 2006), The inducible metabolic capacity of beige fat is a potential target to promote energy expenditure against metabolic diseases (Boss & Farmer, 2012; Tharp & Stahl, 2015). Our mechanistic investigations uncover a novel clock regulation of actin dynamic and MRTF/SRF activity, an inhibitory pathways of beige precursor differentiation. Using gain-and loss-of-function approaches, we further demonstrate that this clock-cytoskeleton-MRTF/SRF regulatory axis impacts energy homeostasis *via* modulation of beige fat activity *in vivo*.

Several key findings from the current study demonstrated the circadian regulation involved in the actin cytoskeleton-MRTF/SRF signaling axis. Transcriptomic profiling in mesenchymal beige precursor stem cells revealed Bmal1 transcriptional regulation of actin cytoskeleton and ECM-related processes, with striking changes in F-actin organization and SRF activity as a result of *Bmal1* loss- or gain-of-function. In beige depot within inguinal subcutaneous fat, SRF and MRTF-A displayed diurnal rhythm, while Bmal1 chromatin association of genes in actin cytoskeleton-MRTF/SRF signaling indicated direct circadian transcriptional control of this pathway. Notably, ECM-cytoskeleton-related genes that are MRTF-A/SRF transcriptional targets were enriched in *Bmal1*-deficient beige depot. Our current study focused on Bmal1 functional targets within the actin cytoskeleton-MRTF/SRF pathway. It is possible that additional circadian clock targets within this signaling network could be uncovered through genome-wide ChIP-seq approach.

In response to extracellular stimuli, including growth factors, cytokines, cell-matrix or cell-cell interactions, monomeric actin polymerizes to form F-actin. The consequent reduction in G-actin level enables MRTF release from sequestration, nuclear translocation, and co-activation of SRF target genes (Esnault *et al*., 2014; Miralles *et al*., 2003). MRTF/SRF-mediated transcriptional response downstream of actin dynamics drives cellular behaviors encompassing cell adhesion, migration, growth and differntiaton (Olson & Nordheim, 2010). The circadian control we identified raises an intriguing possibility of time-of-the-day variations of signaling events within the actin-MRTF-SRF cascade, or signal transduction *via* cell-matrix interactions upstream of actin remodeling, Circadian clock may orchestrate temporal coordination within extracellular matrix-actin cytoskeleton-MRTF/SRF signaling network to impact beige adipocyte development.

Serum-driven clock synchronization in the liver involves cyclic actin remodeling and resultant MRTF/SRF activation (Gerber *et al*., 2013). However, transcriptional regulation of actin cytoskeleton organization by the clock circuit has not been explored. Together with the prior observation of MRTF/SRF-induced clock entrainment (Esnault *et al*., 2014; Gerber *et al*., 2013), the current finding of Bmal1 transcriptional control of actin-MRTF/SRF pathway suggests a regulatory feed-back loop between the circadian clock and cytoskeleton-related processes. As intracellular actin dynamic and the MRTF/SRF transcriptional response transduce extracellular niche signals (Olson & Nordheim, 2010), the reciprocal interactions between the circadian clock and actin-MRTF/SRF axis may facilitate cellular entrainment and adaptation to cyclic cues within its extracellular microenvironment. Little is known to date regarding interactions between circadian clock, beige adipocyte extracellular niche, and beige fat development. Interestingly, while MRTF/SRF suppresses beige precursor development, this pathway promotes a fibrogenic fate decision in adipogenic progenitors during development of obesity (Lin *et al*, 2018; McDonald *et al*., 2015). *Bmal1* deficiency in beige adipocytes, *via* its regulation of actin cytoskeleton-MRTF/SRF activity, induced the thermogenic program whereas its overexpression suppressed beigeing with adipocyte hypertrophy. These findings, in aggregate, suggest a possibility that clock modulation of beigeing we observed may involve the actin-MRTF/SRF cascade in determining between thermogenic induction and energy storage *via* adipogenesis in beige precursor cells (Lin *et al*., 2018; McDonald *et al*., 2015).

Circadian clock exerts temporal coordination of metabolic processes, and its disruption predispose to the risk for obesity and insulin resistance (Panda *et al*., 2002; Turek *et al*., 2005). Determining the contribution of adipose depot clocks to whole-body metabolism, due to their overlapping function and developmental origins, remains challenging. We applied Prx1-Cre transgenic to selectively target beige fat clock modulation (Krueger *et al*., 2014; Logan *et al*, 2002; Sanchez-Gurmaches *et al*., 2015), revealing the metabolic impact of the clock-MRTF/SRF axis with *Bmal1* deficiency-induced beigeing promoting energy expenditure while activating this pathway resulting in obesity. Interestingly, *Bmal1* regulation of MRT/SRF was confined to mature adipocytes, despite Prx1-Cre-mediated high efficiency targeting of both progenitor and adipocytes in beige depot (Krueger *et al*., 2014; Sanchez-Gurmaches *et al*., 2015). Mixed cell populations of SVF in addition to adipogenic progenitors may mask Bmal1 regulation, which could be validated through further analysis using enriched beige progenitors. In HFD-induced obesity, MRTF/SRF activity induces a myo-fibroblast fate switch of adipogenic progenitors to promote adipose tissue fibrosis (Lin *et al*., 2018). Notably, mice subjected to clock disruption by rotating shiftwork developed marked adipocyte hypertrophy with extensive fibrosis in subcutaneous and visceral adipose depots (Xiong *et al*., 2021). Investigation of how dysfunction of the clock-MRTF/SRF cascade induced by circadian misalignment impacts thermogenic capacity or fibrotic response may reveal specific circadian etiologies underlying the adverse metabolic consequences. Identification of a clock-actin-MRTF/SRF regulatory axis as an inhibitory pathway of beige adipocyte development adds to our current understanding of mechanistic links between circadian clock disruption and metabolic diseases.

## Materials and Methods

### Animals

Mice were maintained in the City of Hope vivarium under a constant 12:12 light dark cycle, with lights on at 6:00 AM (ZT0). All animal experiments were approved by the Institutional Animal Care & Use Committee (IACUC) of City of Hope according to guidelines. Experimental procedures for mice studies were carried in accordance with the IACUC approval. *Bmal1*^fl/fl^ and *Prx1*-Cre transgenic mice were obtained from the Jackson Laboratory (Logan *et al*., 2002; Storch *et al*, 2007) and used to generate mice with beige fat-selective *Bmal1* ablation. Conditional ROSA-26 *Bmal1* knock-in mice were generated *via* CRISPR-mediated gene targeting by Cyagen Biosciences. The targeted allele for ROSA-26 *Bmal1* knock-in was shown in Supplemental Fig. S5A. Genotyping primer sequences were listed in Supplemental Table 4. Ambient temperature was set at 21°C. Cold acclimation to induce beigeing was performed in 10 to 12-weeks-old mice at 10°C for two weeks using an environmental chamber with programmable temperature control (Power Scientifics). Thermoneutral adaptation were carried out for two weeks at 30°C. Both male and female mice were used as indicated in the experiments.

### C3H10T1/2 culture, generation of stable cell lines and beige differentiation

C3H10T1/2 were obtained from ATCC, maintained and differentiated in DMEM with 10% fetal bovine serum, as previously described (Liu *et al*., 2020; Nam *et al*., 2015b). Stable lines of *Bmal1* knockdown using shRNA or overexpression using *Bmal1* cDNA were generated as previously described (Nam *et al*., 2015b). To induce beige adipogenic differentiation, induction media containing 1.6μM insulin, 1μM dexamethasone, 0.5mM isobutylmethylxanthine (IBMX), uM Rosiglitazone (Rosi) and 1nM T3 was used for 3 days, followed by maintenance medium for 6 days with insulin, Rosi and T3 (Liu *et al*., 2020).

### Immunoblot analysis

20-40 µg of total protein was resolved on SDS-PAGE gels followed by immunoblotting after nitrocellulose membrane transfer. Membranes were developed by chemiluminescence (Supersignal; Pierce Biotechnology). Antibodies used are listed in Supplemental Table 1.

### RNA extraction and RT-qPCR analysis

Trizol (Invitrogen) and RNeasy miniprep kits (Qiagen) were used to isolate total RNA from snap-frozen tissues or cells, respectively. cDNA was generated using q-Script cDNA Supermix kit (Quanta Biosciences) and quantitative PCR was performed in triplicates on 7500 Fast Real-Time PCR system (Applied Biosystems) with Perfecta SYBR Green Supermix (Quanta Biosciences). Relative expression levels were determined using the comparative Ct method with normalization to 36B4 as internal control. PCR primers sequence are listed in Supplemental Table 2.

### Chromatin immunoprecipitation (ChIP)-qPCR and serum shock

Following formaldehyde fixation, C3H10T1/2 cell chromatin was sonicated and purified. Immunoprecipitation was performed using a specific Bmal1 antibody (Abcam AB93806) or control rabbit IgG using Magna ChIP A/G kit (Millipore), as previously described (Chatterjee *et al*., 2013). Real-time PCR was carried out in triplicate using primers for putative Bmal1 binding E or E’-box elements within ±2kb gene regulatory regions identified using TRANSFAC. Primers flanking Bmal1 E-box within *Nr1d1* promoter were used as a positive control, and upstream primers as a negative control. Values were expressed as fold enrichment over IgG control normalized to 1% of input. Chromatin immunoprecipitation (ChIP) primer sequences were listed in Supplemental Table 3. Serum shock synchronization of C3H10T1/2 cells was induced by 20% FBS treatment for 2 hours following serum starvation overnight, as previously described (Chatterjee *et al*., 2013).

### RNA-seq and computational analysis

Total RNA was extracted from adipose tissue using Trizol (Invitrogen) followed by RNeasy column purification (Qiagen). Sequencing libraries were prepared with Kapa RNA mRNA HyperPrep kit (Kapa Biosystems) according to the manufacturer’s protocol. RNA-seq libraries were sequenced on Illumina HiSeq 2500, as described (Xiong *et al*., 2021). RNA-seq sequencing reads were aligned with STAR software to the mouse reference genome mm10 (Dobin *et al*, 2013), and unique reads were quantified using HTSeq-count with GENCODE annotations (Anders *et al*, 2015). The RNA-seq reads were normalized and log-transformed using limma edgeR packages. P-values and the False Discovery Rate (FDR) were calculated from raw counts using DESeq2 (Love *et al*, 2014). Fold-change > 1.5, 50% FPKM > 0.1, unadjusted p-value < 0.05 and FDR < 0.25 were used as cut-off for differentially expressed genes (DEG). Global analysis heatmaps were produced using heatmap.3 and the gplots package in R, using log2(FPKM + 0.1) values. The Pearson Dissimilarity (1-Pearson correlation coefficient) was used as the distance metric for hierarchical clustering of rows and columns. Expression was centered so each gene had a mean of 0, and centered expression was capped at -3 and 3, for clustering and visualization. This RNA-seq dataset was deposited in NCBI GEO GSE183000.

### Primary preadipocyte isolation and beige differentiation

The stromal vascular fraction containing preadipocytes were isolated from subcutaneous or gonadal fat pads, as described (Chatterjee *et al*., 2013). Briefly, dissected fat pads were cut into small pieces and digested in 0.1% collagenase Type 1 in DMEM with 0.8% BSA at 37°C in a horizontal shaker for 60 minutes. The digestion mixture was passed through Nylon mesh and centrifuged to collect the top adipocyte layer and the pellet containing the stromal vascular fraction with preadipocytes. Preadipocytes were cultured in F12/DMEM supplemented with bFGF (2.5 ng/ml), expanded for two passages and subjected to differentiation in collagen type I-coated 6-well plates at 90% confluency. Beige adipocyte differentiation was induced for 2 days in induction medium (10% FBS, 1.6 μM insulin, 1 μM dexamethasone, 0.5 mM IBMX, 0.5uM Rosi and 1nM T3), before switching to maintenance medium for 4 days (insulin, Rosi and T3). Jasplakinolide (0.1μM) and CCG-1423 (5 μM) were obtained from Cayman Chemicals and added for the entire differentiation time course.

### Oil-red-O, Bodipy and Mitotracker staining

These staining procedures were performed as previously described (Nam *et al*., 2015b). Briefly, for Oil-red-O staining, cells were fixed using 10% formalin and 0.5% Oil-red-O solution was incubated for 1 hour. Bodipy 493/503 was used at 1mg/L together with DAPI for 15 minutes, following 4% paraformaldehyde fixation and permeabilization with 0.2% triton-X100. Mitotracker Deep Red FM was applied at 100 nM to differentiated adipocytes and incubated for 30 minutes prior to fixation (Liu *et al*., 2020; Nam *et al*., 2015a).

### Hematoxylin/eosin histology and adipocyte size distribution

Adipose tissues were fixed with 10% neutral-buffered formalin for 72 hours prior to embedding. 10μm paraffin sections were processed for hematoxylin and eosin staining. Adipocyte size area was measured by outlining the adipocytes, and the average of five representative 10X fields from each mouse were plotted for cross section area distribution, as described (Xiong *et al*., 2021).

### Glucose tolerance test, insulin tolerance test and insulin signaling

The glucose tolerance test (GTT) and insulin tolerance test (ITT) were performed as described (Yin *et al*, 2020). Mice were fasted overnight prior to GTT. The indicated dose of glucose used was administered by i.p. injection. ITT were performed following 4 hours of fasting with doses specified. Insulin signaling in adipose tissues was determined at baseline in mice fasted for 4 hours. For insulin stimulation, tissues were collected 20 minutes after intraperitoneal injection of 0.5U/Kg insulin.

### Hyperinsuinemic-euglycemic clamp

The low-dose hyperinsuinemic-euglycemic clamp studies were performed as described (Yin *et al*., 2020). Mice were cannulated through the right jugular vein, allowed 4 days of recovery and studied in a conscious state. Briefly, a priming dose (10 μCi) and a constant intravenous dose (0.1 μCi/min) of ^3^H-glucose were infused through the venous canula. Basal glucose production was assessed after 50 minutes of glucose infusion. Mice were then primed with a bolus insulin followed by continuous infusion at 3 mU/kg/min. During this time, glucose was simultaneously infused to maintain a steady level at 100-140 mg/dL. For the analysis of insulin-stimulated glucose tissue uptake, 2-deoxy-D-[1-^14^C] glucose was administered as a bolus 45 minutes prior to completion of the clamp study. Individual tissues were snap-frozen and the ^14^C-glucose uptake was assayed by scintillation counting and calculated from ^14^C-glucose plasma profile fitted with a double exponential curve and tissue content.

### Indirect calorimetry by Comprehensive Laboratory Animal Monitoring System (CLAMS)

Mice were single housed and acclimated in metabolic cages using the Comprehensive Laboratory Animal Monitoring System (Columbus Instruments), with ad libitum food and water with controlled lighting for 2 days prior to metabolic recording. Metabolic parameters, including oxygen consumption, CO2 production, respiratory exchange ratio, ambulatory activity, and food intake were recorded for 5 consecutive days, as described (Yin *et al*., 2020).

### Statistical analysis

Data was expressed as mean ± SEM. Statistical analysis was performed using GraphPad Prism. The differences between groups were determined by unpaired two-tailed Student’s t test, or one-way ANOVA with post-hoc pairwise t tests with Bonferroni correction. A minimum of three biological replicates were used to perform statistical analysis. P values ≤0.05 were considered statistically significant.

## Non-standard Abbreviations

MRTF: Myocardin-related transcription factor
Bmal1: Brain and Muscle Arnt-like Protein 1
CLOCK: Circadian Locomotor Output Cycles Kaput
ROR: RAR-related Orphan Receptor
Prx1: Paired-box Related Homeobox 1

## Authorship contributions

Xuekai Xiong, Weini Li and Ruya Liu: data curation, formal analysis, methodology and investigation; Vijay Yechoor: data curation, manuscript review and editing, and funding acquisition; Ke Ma: data curation, formal analysis, project administration, manuscript writing and editing, and funding acquisition.

## Acknowledgements

We thank the Shared Resources Core Facility of City of Hope for their expert technical assistance on RNA-Seq analyses. K. M. is a faculty member supported by the NCI-designated Comprehensive Cancer Center at the City of Hope National Cancer Center. We also thank the Metabolic Phenotyping Core Laboratory at Baylor College of Medicine for their expert technical support with insulin clamp studies. This project was supported by grants from National Institute of Health 1R01DK112794, and American Heart Association 17GRNT33370012 to K. M.; and National Institute of Health grant DK097160-01 and 1R01DK130499 to V. Y.

